# Monitoring and maturation of cardiomyocyte differentiated from human induced pluripotent stem cells using a multi-functional tissue engineering platform

**DOI:** 10.1101/2023.05.15.540892

**Authors:** A-Ri Kim, Sajal Shrivastava, Han-Byeol Lee, Nae-Eung Lee

## Abstract

**Background:** The use of human induced pluripotent stem cells (hiPSCs) is attracting attention as a potential personalized therapy for damaged myocardium, as they are accessible and compatible with the human body. The hiPSCs can be effectively differentiated into cardiomyocytes (CMs), which are essential for myocardial regeneration. However, current approaches have been unable to fully replicate the electro-mechanical functions of natural CMs.

**Methods:** Although several methods have been used to stimulate hiPSC-CMs for maturation, the ability to integrate multiple stimuli that influence myocardial cell function has been limited. To address this challenge, we have developed a multi-functional tissue engineering system for CMs that is based on a stretchable multielectrode array (SMEA). The SMEA is housed in a mini-incubator and is stretchable, durable and autoclavable.

**Results:** The system can independently control co-stimulation parameters of electrical and mechanical stimuli simultaneously. Furthermore, the system can monitor the status of the cells throughout their proliferation and differentiation of hiPSCs, as well as the stimulation of hiPSC-CMs, through electrochemical cell impedance spectroscopy. By applying co-stimulation, we have observed the enhanced maturation of hiPSC-CMs.

**Conclusions:** Our innovative system holds great potential as a tool for improving the culture and engineering of electrogenic cells with enhanced maturity.

## 1. Introduction

The mechanical contraction events of cardiac muscle are triggered by electrical stimulation that provokes a calcium-dependent excitation-contraction coupling. Dysregulation of mechanical contraction and voltage-dependent ion channels results in various life-threatening conditions including arrhythmias and heart failure. Recent developments in stem cell (SC)-based approaches have created promising results in regenerative cardiac therapy that targets the repair of irreversibly damaged cardiac tissues. For example, novel personalized treatment strategies using reprogrammed human induced pluripotent SCs (hiPSCs) that can be differentiated into a variety of cell lines including cardiomyocytes (CMs) are attracting a great deal of attention due to their easy accessibility and excellent histocompatibility^1–4^. However, cell-based regeneration therapies based on the heart implantation of functional cardiac grafts have very limited success due to the extremely poor survival rate of the cardiac tissue construct^5^. Furthermore, *in vitro* models have been used recently to study regenerative cardiac therapies such as organoids, spheroids and organ-on-a-chip based on hiPSCs derived from patients as personalized therapeutics to treat heart diseases^6–8^. However, current therapies and models have yet to fully replicate the electro-mechanical function of the myocardium. The limited success of these models is due to their inability to combine various cell stimulations that impact the function of CMs, which ultimately control the microenvironment of the myocardium. Therefore, achieving the electro-mechanical maturation of CMs is a critical functional parameter that determines the effectiveness of cardiac regenerative therapy^9–17^.

For this purpose, various attempts have been undertaken to develop active and passive devices for electrical and mechanical stimulations to induce functional maturity in hiPSC-derived CMs (hiPSC-CMs) to mimic adult cardiac tissue constructs. For example, mechanical stimulations have been achieved by several methods including scaffolds with varying material stiffness, compression, flow-driven shear stress, mechanical stretching and three-dimensional (3D) nanotopography^8, 18–23^. Furthermore, various electrical stimulation platforms have been developed to induce excitation-contraction coupling and synchronous spontaneous beating in CMs^24–27^. Although the experimental results of mechanical and electrical co-stimulation for CMs derived from hiPSCs also have shown promising outcomes^20, 23, 24, 28, 29^, there are still several issues that need to be addressed. One of the main challenges is in achieving reproducibility and consistency of the results. The system requires optimization of various parameters, such as the mechanical and electrical stimulation conditions, to achieve optimal results during reliable long-term culture and co-stimulation.

Herein, we propose an integrated highly durable multi-functional tissue engineering system which can co-stimulate the hiPSC-CMs with electrical and mechanical cues simultaneously for enhanced maturation and at the same time monitor different stages of proliferation and differentiation of hiPSCs and co-stimulations of hiPSC-CMs. The system integrated within a mini-incubator provides the capability of culture and differentiation of hiPSCs and also of applying mechanical and electrical stimuli simultaneously for the maturation of hiPSC-CMs, which is enabled by forming a stretchable, mechanically durable and autoclavable multielectrode array (SMEA). The monolithic integration of electrical and mechanical stimulation components and the dual function of the SMEA allow a simple structure with a small footprint. The SMEA. device formed on a 3D patterned stretchable substrate with the capability of stress absorption for the electrode region and electrical tracks is used for co-stimulation and monitoring purposes. The combination of different electrical and mechanical stimulation conditions in various ranges has enabled us to tune the co-stimulation conditions that affect the maturity of the hiPSC-CMs. The hiPSC-CMs subject to co-stimulations were found to be matured more effectively compared to those subjected to either mechanical or electrical stimulation only. The state of the hiPSC-CMs contained in the well could be monitored through optical microscopy and electrochemical cell impedance spectroscopy (ECIS). The proposed approach has great potential for generating matured cardiac cells that can be used in a variety of fields such as cell therapy, drug screening, disease modelling and personalized medicine.

## 2. Methods

### Construction of a multi-functional tissue engineering system

#### Stretchable 3D micro-patterned substrate

To produce a stretchable 3D micro-patterned PDMS, a positive PR (ma-P 1275HV, Micro Resist Technology) is first patterned on a glass substrate (Taewon Scientific) by a double photolithography method, and then a PDMS (Sylgard A+B, 10:1, Dow Corning) and a PUA mould are produced in order by a soft lithography method^30^. Elastomer PDMS was poured into the PUA master mould to a thickness of about 500 μm, and then cured for 2 hours naturally for even flatness and baked in an oven at 80 °C. The stretchable 3D micro-patterned PDMS substrate was removed from the master mould, cut into 1.5 cm x 3.5 cm sections and prepared as a substrate.

#### Fabrication of the SMEA device in a cell culture well

After attaching a stencil mask manufactured to match the shape of the electrode to the 3D micro-patterned PDMS substrate, a 3-nm-thick Al_2_O_3_ adhesion promotion layer was deposited by atomic layer deposition. Then, e-beam evaporation was used to deposit a 7-nm-thick Ti glue layer and a 63-nm-thick Au electrode layer using a shadow mask. Six holes for the opening of the electrode tip areas were punched into the encapsulation layer with a diameter of 350 μm using a micro-scale punching needle to enable electrical signal transmission and signal reading. The encapsulation layer and 3D micro-patterned PDMS substrate were aligned and treated with oxygen plasma to create strong Si-O-Si bonds between the two layers. Oxygen plasma (O_2_ flow rate of 1500 sccm and N_2_ flow rate of 500 sccm) in a microwave plasma reactor was applied for 20 s for this purpose. For strong adhesion of the two layers, the assembly was kept under a pressure of 600 psi in a hot-pressing machine at 90 °C. For the cell culture, a cylindrically shaped PDMS well with an opening is attached to the top of the encapsulation layer. For PDMS well fabrication, PDMS is poured into a 48-well plate (Corning), a 10-mL serological tube was inserted to close the hole, and then it was cured in an 80 °C oven for 2 h. Before the use of the device, after washing sufficiently with ethanol, the PDMS was subjected to oxygen plasma treatment (O_2_ flow rate of 1500 sccm and N_2_ flow rate of 500 sccm) for 30 s to create a hydrophilic surface.

#### Set-up of the multi-functional tissue engineering system

Inside a commercial mini-incubator (Live Cell Instruments) with temperature control, medium supply and O_2_/CO_2_ gas supply functions, a customized PCB capable of applying electrical stimulation and reading the signal from the electrodes was installed, and a motor with cyclic uniaxial extension was installed on the right side of the incubator. This system allows the user to provide mechanical and electrical stimuli independently at desired values. The system reduces the damage to cells during stimulation and provided the cells with an appropriate temperature, humidity, and carbon dioxide concentration for long-term culture and monitoring on the SMEA device. **Supplementary Video 7** displays a multi-functional tissue engineering system that provides both electrical and mechanical co-stimulation.

### hiPSC line and hiPSC seeding on SMEA

#### Stem cell lines

The hiPSCs used in the experiment were the cells from cord blood and two cell lines purchased from two providers (Thermo Fisher Scientific and Accegen). They were cultured on 100-mm Petri dishes coated with human VTN-N truncated recombinant protein (Gibco, Thermo Fisher Scientific). The hiPSCs obtained from Thermo Fisher Scientific were maintained in Essential 8 Medium (Gibco, Thermo Fisher Scientific) and the hiPSCs obtained from Accegen were maintained with mTeSR (STEMCELL Technologies) at 37 °C with 95% air and 5% CO_2_. The hiPSCs were subcultured in the devices at 70-80% confluence using TrypLE Express Enzyme (Gibco, Thermo Fisher Scientific). First, we coated the culture wells with 150 μl of VTN-N mixed with Dulbecco’s phosphate-buffered saline (DPBS) solution at a concentration of 5 μg/mL and incubated at room temperature for 1 hour. After removing the VTN-N at d 1, hiPSCs were seeded on the devices at a confluency of 1×10^5^/mL (total of 4×10^4^ cells in the culture well).

#### Proliferation and differentiation

The hiPSC medium was changed every day for 3 days and the seeded hiPSCs were allowed to proliferate until a confluency of 80% was reached on the device. The seeded hiPSCs were differentiated using a differentiation kit (PSC CM differentiation kit, Gibco, Thermo Fisher Scientific). After removing the hiPSC culture medium at d 3, we added CM differentiation medium A and incubated them for 2 days and replaced it with CM differentiation medium B solution at d 5. From d 7, the medium was changed to CM maintenance medium once every two days. From d 3 to d12, the hiPSCs were differentiated into hiPSC-CMs.

### Electro-mechanical co-stimulation

The stimuli of uniaxial cyclic stretching at 1 Hz and a biphasic electrical stimulation 7 with a duration of 1 ms were given every day for 1 hour for a total of 5 days. The electrical stimulation was given with a biphasic waveform generated from a function generator (AFG3102, Tektronix), which is applied to the SMEA device with the culture well through the PCB inside the mini-incubator. The applied electrical stimulation is transmitted to the electrodes across the opposite sides. The stimulation signal was monitored through an electrometer (SMU 2612, Keithley Instruments). The motor installed on the right side of the mini-incubator enables uniaxial cyclic stretching. By adjusting the sample fixing screws inside the incubator to set the spacing of the samples, the extension level can be set and the desired speed can be adjusted through sensors inside the motor box.

### Viability assay

A viability assay was carried out using a viability assay kit (CELLOMAX^™^, Precaregene). The product uses WST-8, a water-soluble tetrazolium salt that reacts with dehydrogenase in the mitochondrial electron transport system of metabolically active living cells, to produce orange-coloured water-soluble formazans, and thus has a linear correlation with the number of living cells. Through viability assay, we compared how electro-mechanical co-stimulation affects the relative viability of hiPSC-CMs that have undergone proliferation as hiPSCs, differentiation from hiPSCs and maturation by co-stimulation. For the test, 40 μL of the solution from the kit was added to the inside of the well of each device and was reacted in an incubator for 4 hours. Then, within 1 minute it was transferred to a 96-well plate and the absorbance was measured at 450 nm using a plate reader.

### Impedance spectroscopy (ECIS) measurements

To investigate the durability and functionality of the SMEA device with and without cells and the status of the cells on the device, ECIS measurements were performed using a potentiostat (VSP, Bio-Logic). The proliferation of hiPSCs as well as differentiation and co-stimulation of hiPSC-CMs on the device were also monitored by measuring the impedance. The counter electrode and a reference electrode (mf-2052 Ag/AgCl, Bioanalytical Systems, Inc.) were used. We measured the impedance spectra with a commercial counter and a reference electrode, which were immersed in a cell culture medium in a cell culture well installed on the SMEA device.

### Immunocytochemistry analyses

After removing the CM maintenance medium, 200 μL of fixative solution (4% formaldehyde in DPBS, Life Technologies) was added and incubated for 15 minutes at room temperature. After removing the fixative solution, 200 μL of permeabilization solution (1% Saponin in DPBS, Life Technologies) was added. After 15 minutes of incubation at room temperature, it was changed to 100 μL of blocking solution (3% BSA in DPBS, Life Technologies) and incubated for 30 minutes at room temperature. Then, 0.1 μL of primary antibodies anti-NKX2-5 (host: rabbit) and anti-TNNT2 (host: mouse) were added directly into the culture well containing 100 μL of blocking solution and incubated for 4 hours at room temperature. All liquids were removed and the device was gently washed 3 times with a wash buffer (Life Technologies). The device was incubated for 1 hour at room temperature in 0.4 μL of Alexa Fluor 594 donkey anti-rabbit (for anti-NKX2-5, Life Technologies) and Alexa Fluor 488 donkey anti-mouse (for anti-TNNT2, Life Technologies) secondary antibodies diluted with 100 μL of blocking solution and then gently washed 3 times with a wash buffer. One drop of DAPI (NucBlue Fixed Cell Stain, Life Technologies) in 5 mL of the wash buffer was added and incubated for 5 minutes.

### mRNA quantification by qRT-PCR

#### RNA extraction

Total RNA was extracted from the cells by using TRIzol reagent (Invitrogen). First, the cells were homogenized with TRIzol reagent and 0.2 mL of chloroform was added to each tube. After vigorously shaking the tube by hand for 15 s, the tube was left to stand at room temperature for 3 minutes. For phase separation, the samples were centrifuged at 13,000 rpm for 15 minutes at 4 °C and the upper aqueous layer was transferred to a new tube. An equal volume of isopropyl alcohol was added and incubated for 10 minutes at room temperature. To precipitate the RNA, the tube was centrifuged at 13,000 rpm for 10 minutes at 4 °C. The supernatant was removed, and the RNA pellet was washed with 1 mL of 70% ethanol. The samples were briefly vortexed and centrifuged at 13,000 rpm for 5 minutes at 4 °C. The supernatant was removed, and the RNA pellet was resuspended in RNase-free water.

#### Measurement of RNA yields, purity and integrity

RNA yield was measured based on the absorbance at 160 nm, and the A_260:280_ and the A_260:230_ ratios were used to assess the purity of RNA using a NanoDrop 2000 Spectrophotometer (Thermo Fisher Scientific Inc.). The RNA integrity number (RIN) was determined by using a high-sensitivity RNA Screen Tape Kit following the manufacturer’s protocol on an Agilent Tape station 42000.

#### mRNA quantification by qRT-PCR

With total RNA, reverse transcription was performed using SuperScript II RNase (Invitrogen) according to the manufacturer’s instructions. The cDNA of the mRNA was amplified using the following primer pairs: MYH6 Forward: 5‘-AGAGTCGGTGAAGGGCATGA-3’, Reverse: 5‘-AGCCGCAGCAGGTTCTTTTT-3’; MYH7 Forward: 5‘-GAGCCTCCAGAGCTTGTTGA-3‘, Reverse: 5‘-ACGATGGCGATGTTCTCCTT-3‘; CACNA1C Forward: 5‘-ATGACGAAAATCGGCAACTG-3’, Reverse: 5‘-GGAAACCCCTCTTCGGAGAT-3‘; CTTCTGCATACGATCAGCAA-3’; GAPDH Forward: 5‘-CGAGATCCCTCCAAAATCAA-3’, Reverse: 5‘-CCTTCTCCATGGTGGTGAA-3’. Real-time PCR was performed on a StepOnePlus^TM^ Real-Time PCR System (Applied Biosystems) using an SYBR Green PCR Kit (Applied Biosystems), according to the manufacturer’s instructions. Thermal cycling conditions were 95 °C for 10 minutes followed by 40 cycles of 95 °C for 15 s and 1 minute at optimal T_m_ (59 °C). The data were analysed using StepOne software v2.2.2 (Applied Biosystems). The expression levels of each mRNA were normalized to an endogenous control GAPDH and were calculated using the 2^-ΔΔCt^ method.

### Calcium transient assay

Fluorescent indicators of Ca^2+^ were used to measure calcium signalling stimulated by agonists and inhibited by antagonists via G protein-coupled receptors (GPCRs), which represent a significant and active target group in drug discovery. Calcium assay buffer (1 mL, Fluo-4 Direct calcium assay buffer, Invitrogen) was mixed with 77 mg of water-soluble probenecid (Invitrogen) to obtain a 250 mM solution of probenecid and the mixture was agitated in a vortex until dissolved. To make a 2× calcium reagent loading solution, 200 μL of 250 mM probenecid stock solution was mixed with 10 mL of Fluo-4 Direct calcium assay buffer. The 2× calcium regent loading solution (200 μl) was added directly into the cell culture well containing 200 μL of cell culture medium. The devices were incubated at 37 °C for 30 minutes and at room temperature for 30 minutes. Confocal images of the cells on the 3D stretchable micro-patterned cell culture device were taken directly. To obtain a calcium transient intensity video, the cell culture medium and calcium reagent loading solution were removed and the cell culture well was cut off. The cell side of the devices was turned down for imaging using an inverted microscope (IX71, Olympus) to allow a path of light at an excitation wavelength of 494 nm and an emission wavelength of 516 nm of the wavelength to pass through.

### Measurement of membrane potential

Various physiological parameters such as cell signalling and muscle contraction, are significantly affected by changes in the electrical potential across the membrane. To prepare a membrane-sensitive loading solution, 10 μL of membrane-sensitive dye (Fluovolt dye, Invitrogen) was mixed with 100 μl of 100× Pluronic™ surfactant polyols (PowerLoad Concentrate, Invitrogen). Then, 10 mL of 20 mM glucose in Hank’s balanced salt solution (HBSS) (Invitrogen, Thermo Fisher Scientific) was added and mixed. The devices containing cell culture medium were washed twice with HBSS. Membrane-sensitive loading solution was added and cells were incubated at room temperature for 30 minutes. Membrane-sensitive loading solution was removed and devices were washed twice with HBSS. To take confocal images, 100 μl of 20 mM glucose stock in HBSS was added to the devices.

### Optical and fluorescence microscopy

Device stability and durability before and after mechanical and electrical stimulation was evaluated using field-emission scanning electron microscopy (FE-SEM, JEO JSM-6500F) and optical microscopy (TH3, Olympus). Images of cells proliferated on the devices were taken by an inverted optical microscope (AE31, MOTIC). Imaging of ICC, calcium transient and membrane potential was carried out using confocal microscopy (FV3000, Olympus). The calcium transient video was examined using a fluorescence microscope (IX71, Olympus).

### Analysis of ICC, calcium transient and membrane potential images

An ImageJ software program (FIJI) was used for the analysis of fluorescence images, including ICC, cell alignment, calcium transient and membrane potential images. We analysed the peaks of calcium transient with a plug-in called Spiky in ImageJ. Statistical analysis for the calcium transient images was performed using GraphPad Prism 9, with one-way analysis (ANOVA) conducted between groups and post hoc Tukey test used for comparisons between two groups. Statistically significant p values were less than 0.05. Data were presented as means ± standard error of the mean.

## 3. Results

### Concept of multi-functional tissue engineering system

A tissue engineering system that possesses the co-stimulation capability of controlled simultaneous electrical pulsing (E) and mechanical strain (ε) to hiPSC-CMs was designed and fabricated (**Fig. 1a**). The developed system was also capable of monitoring the stages of cell culture using ECIS. After stimulation or monitoring, the cells were investigated additionally through immunocytochemistry (ICC), gene expression analysis, calcium transient assay and membrane potential measurements. The multi-functional tissue engineering system is primarily composed of an SMEA device on a 3D micro-patterned stretchable polydimethylsiloxane (PDMS) substrate with a PDMS well, which can be loaded into a commercial culture incubator for long-term culture and a mini-incubator housed with the SMEA device (**Fig. 1b**). It was designed in such a way that it can be mounted on an optical or confocal microscope for the observation of cells. The SMEA device was fabricated by forming six Au electrodes on the 3D micro-patterned stretchable PDMS substrate and encapsulated with a PDMS layer. The mini-incubator was tailored to maintain optimal levels of CO_2_, humidity and temperature during co-stimulation and ECIS measurements. A specifically designed printed circuit board (PCB) enabled an electrical connection between control electronics and the SMEA for electrical stimulation and cyclic uniaxial extension for mechanical stimulation. During co-stimulation, the status of electrical stimulation on the tissue engineering system was monitored continuously using an electrometer. For the optimal operation of the system, multiple combinations of simultaneous electrical and mechanical cues were tested to understand their effects on cell maturation. **Supplementary Fig. 1** shows images of the components in the multi-functional tissue engineering system and the connection between them. The main components include the SMEA device, control components for co-simulations, the modified mini-incubator for cell culture, the monitoring and imaging equipment for the cells on the SMEA device, maintenance components of the culture environment and peristaltic pump for the supply of the culture media to the culture well in the SMEA device during stimulation and monitoring.

**Figure 1.**
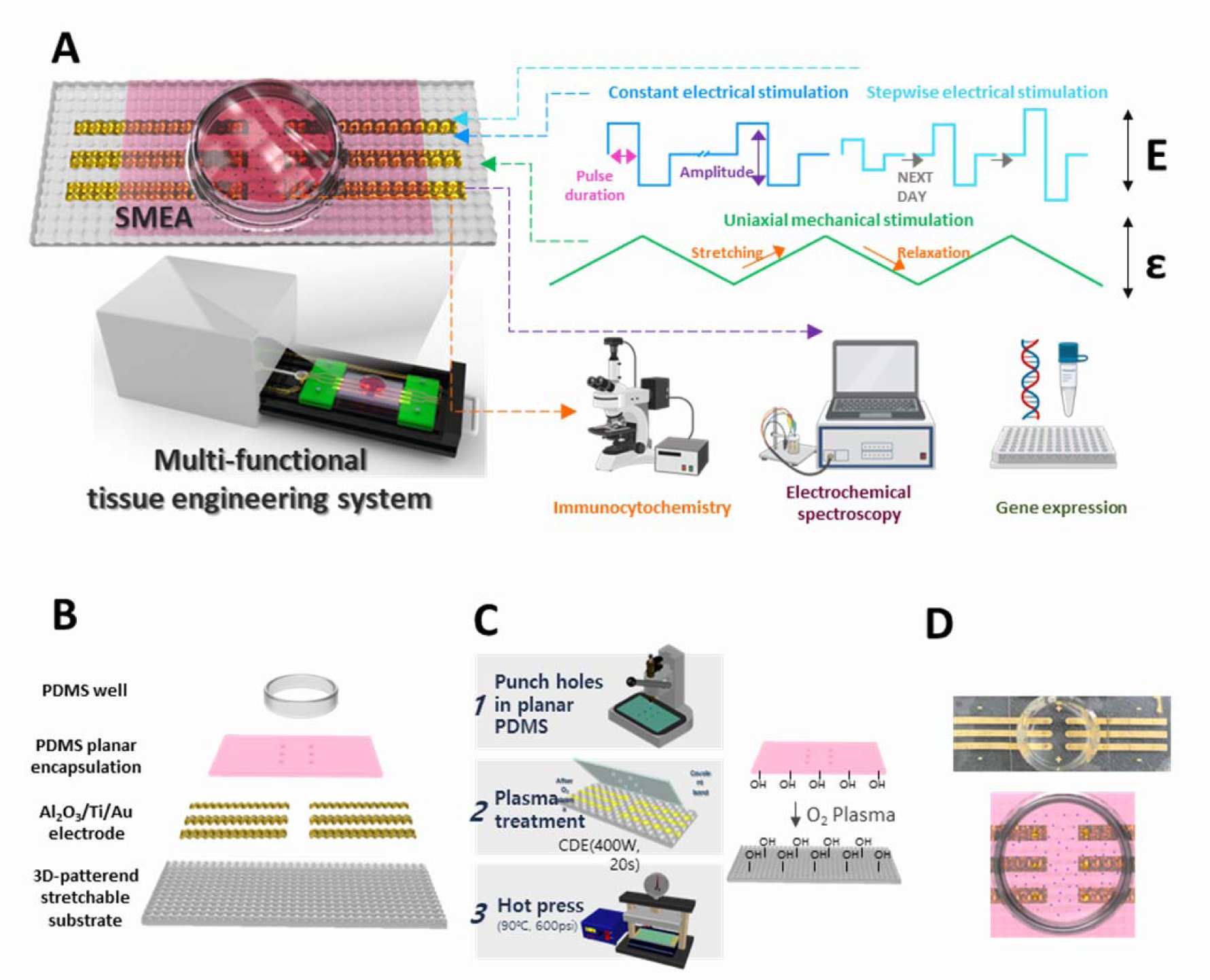
Schematic and fabrication of multi-functional tissue engineering system. **A**, Schematic of the multi-functional tissue engineering system which is composed of a stretchable multi-electrode array (SMEA) device for culture, simultaneous stimulation of electrical pulsing (E) and uniaxial extensional straining (ε), and monitoring of cells and a mini-incubator housed with the SMEA device. The cells can be monitored with electrochemical cell impedance spectroscopy (ECIS) and also analyzed though immunocytochemistry as an end-point cell analysis. The stimulated cells were harvested and used for further gene expression analysis. **B**, An exploded schematic view of the SMEA device with a culture well. **C**, Encapsulation of the Au electrodes with an encapsulation layer. **D**, Top view of a fabricated SMEA (top) and schematic of the SMEA with cell loading (bottom)

### Fabrication of highly durable SMEA device

A highly durable SMEA device was required with the capability of co-stimulating hiPSC-CMs over an extended time under the influence of the culture medium. The key factor for the successful co-stimulation and monitoring of the SMEA operation was the structural engineering of the PDMS substrate to avoid electrode damage during repetitive mechanical stimulation while applying electrical pulsing in the liquid medium. For this purpose, the 3D micro-patterned substrate, which comprised curvilinearly connected valleys and bumps with no planar region to enable stress absorption on the electrode region under cyclic stretching, was used to allow uniaxial stretchability of the electrodes of up to 30% without significant damage^30, 31^. The transparency of the PDMS with relatively low background noise allows optical imaging. The stretchable 3D micro-patterned substrate was moulded from a UV-curable poly(urethane acryl) (PUA) mould replicated from a master mould fabricated on a Si wafer using a double lithography method^30^.

For the fabrication of Au electrodes, three layers of Al_2_O_3_, Ti and Au were deposited in sequence onto the 3D micro-patterned substrate and then encapsulated (**Fig. 1c**). The substrate formed a six Au electrode array that was encapsulated by a PDMS layer except for the electrode tips where cells adhere and are subject to mechanical straining during cyclic stretching. To contact the cells on the stretchable electrode array, six holes were punched in the encapsulation layer to expose the tip area of each electrode for electrical signal transmission and signal reading. The encapsulated planar area of the device where most cells existed covered 96% of the total substrate surface area. The encapsulation layer also provides a neutral plane for the electrodes and therefore reduces the stress applied on the track of the electrode^32^. The encapsulation layer and 3D micro-patterned substrate were treated with microwave oxygen plasma and hot-pressed to create a Si-O-Si bond between the two layers for strong bonding^33^. Finally, a PDMS well for the containment of cells and culture medium was attached. The picture and schematic image of the fully assembled SMEA with the PDMS well attached is shown in **Fig. 1d**.

### Evaluation of durability and performance of SMEA device

To evaluate the durability of the SMEA device, we carried out 40,000 cycles of mechanical stretching at 20% ε and 1 Hz frequency while the device was immersed in cell culture medium inside the well. **Fig. 2a** shows the field-emission scanning electron microscopy (FE-SEM) images of the device after cyclic stretching. Fig. 2a(I) shows the top view of the well electrode area. A cross-section FE-SEM image of the gap area between the 3D micro-patterned PDMS substrate shows that the PDMS encapsulation layer is stable and firmly attached without delamination at the interface (Fig. 2a(II)). The tiled cross-sectional and enlarged cross-sectional FE-SEM images of the electrode area shown in Fig. 2a(III) and 2(IV), respectively, indicate no observable damage after cyclic stretching of the SMEA in culture media.

**Figure 2.**
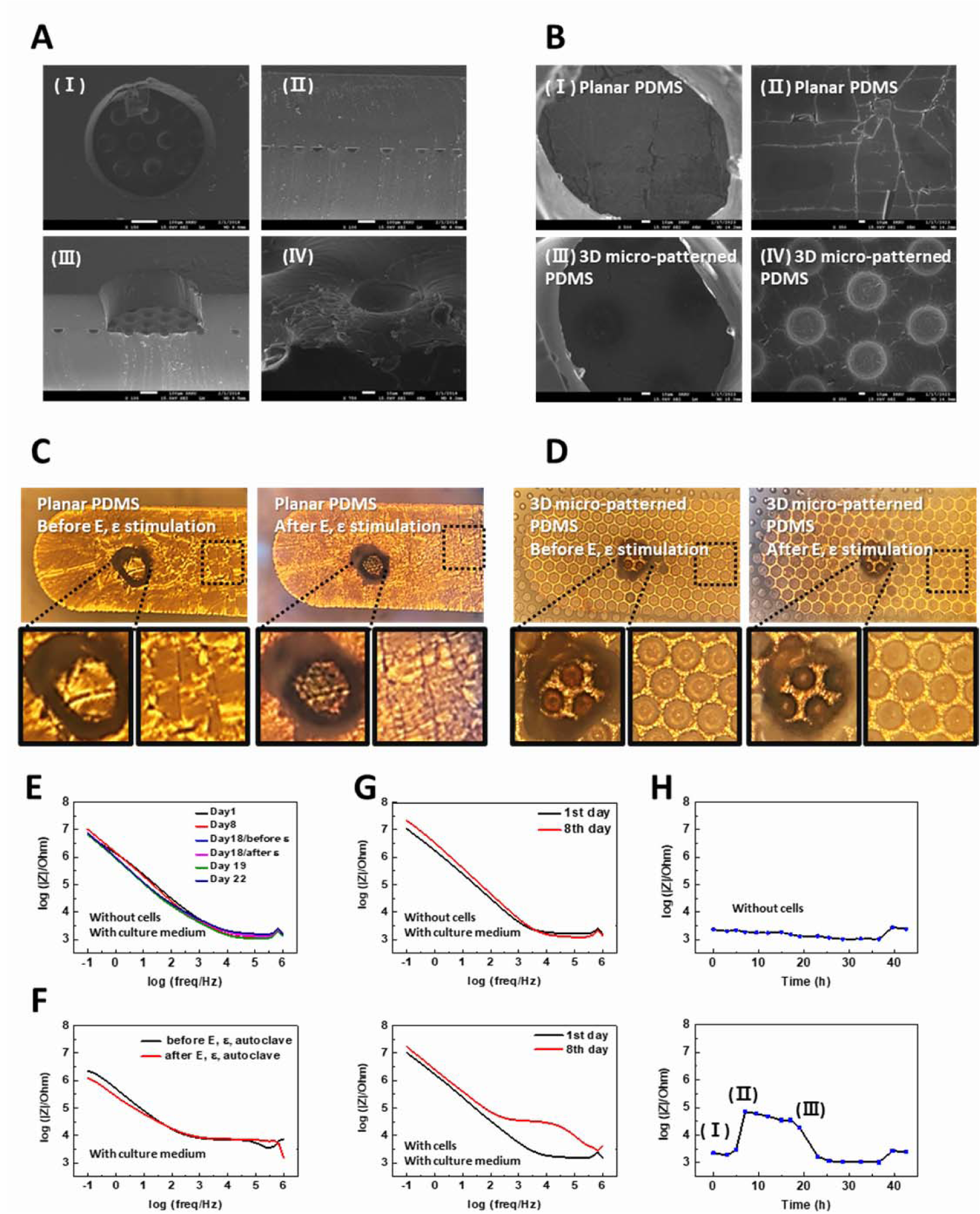
Evaluation of durability and functionality of the SMEA device. **A** (I) Top view, (II) cross-sectional, (III) tilted and (IV) magnified top view FE-SEM images of the Au electrode areas of the device after cyclic stretching 40,000 times in culture medium at 20% of strain (ε). Scale bars I, II, III: 100 μm, IV: 10 μm. **B**, (I), (III) top view and (II), (IV) magnified top view FE-SEM images of the electrode area on planar ((I) and (II)) and 3D micro-patterned substrate ((III) and (IV), respectively. **C**, **D**, Optical images of the device on planar and 3D-micro-patterned PDMS substrate before and after electrical and mechanical co-stimulation in culture medium, respectively. **E**, Impedance spectra of a bare device before and after cyclic stretching (10,000 times at 10% of ε) measured at d 18 while monitoring from d 1 to d 22. **F**, Impedance spectra of a bare device before and after co-stimulation (50,000 stretching cycles at 10% of ε and 5 V/cm of E stimulation) and followed autoclaving. **G**, Impedance spectra obtained from the device without (upper panel) and with (lower panel) H9C2 (rat myoblast) cells at d 1 and d 8. **H**, Impedance (log (|Z|/Ohm) values during long-term monitoring of the device without (upper panel) and with (lower panel) H9C2 cells for 2 d at 10kHz. With the cells on SMEAs, impedance change due to initial seeding (I), cell-electrode adhesion (II) and detachment (III) can be observed.

For further investigation of device durability, the morphological changes of Au multi-electrodes formed on the planar stretchable and 3D micro-patterned PDMS substrates were observed and compared after cyclically stretching the device 50,000 times at 10% ε with a frequency of 1 Hz while applying the biphasic electrical pulsing between two electrodes at E of 5 V/cm and the frequency of 1 Hz in the culture medium. The top view in FE-SEM images of the device after co-stimulation in the liquid medium shows long and large cracks from the electrode fabricated on the planar PDMS substrate (**Fig. 2b**(I) and **2b**(II)) but a small number of short cracks on the 3D micro-patterned PDMS substrate (**Fig. 2b**(III) and **2b**(IV)). The optical microscope images of the cyclically stretched samples indicate that many cracks already exist in the electrodes on the planar PDMS substrate even before the co-stimulation (**Fig. 2c**, left panel) and the crack density was increased after co-stimulation (**Fig. 2c**, right panel) whereas the electrodes formed on the 3D micro-patterned PDMS substrate have some wrinkles but cracks could not be observed before the co-stimulation (**Fig. 2d**, left panel) and after co-stimulation (**Fig. 2d**, right panel).

To verify the electrical stability of the electrodes, the device without cell loading was incubated with cell culture medium for 22 days, and cyclic stretching of 10,000 times at 20% of ε with a frequency of 1 Hz was applied d 18 while ECIS measurements were carried out from d 1 to d 22. ECIS measurements were carried out on the same device during the incubation period as well as before and after stimulation of the device to see any change in the electrical properties. The results shown in **Fig. 2e** show that there is no significant change in the measured impedance spectra indicating excellent electrical stability of the electrode array after repetitive mechanical straining and long-term storage in the culture medium. The SMEA seeded with H9C2 cells on d 8 were also cyclically stretched by 10,000 times at the ε of 20% on d18 and the impedance spectrum was measured (**Supplementary Fig. 2**). These results indicate that the device without cell seeding resulted in no significant change in the impedance whereas the device showed a significant change upon cell seeding and mechanical stimulation presumably due to the movement and rearrangement of the cells. No significant change in the impedance before and after electro-mechanical co-stimulation even when the device was cyclically stretched by 50,000 times at the ε of 10% with the frequency of 1 Hz while the biphasic electrical pulsing with 5 V/cm between two electrodes at the frequency of 1 Hz was applied in culture medium (**Supplementary Fig. 3**). After electro-mechanical co-stimulation, the device was rinsed with 70% ethanol and distilled water, and then autoclaved at an elevated temperature of 121 °C and high pressure of 26 psi for sterilization. A comparison of the data in **Fig. 2f** showed that the impedance values (log (|Z|/Ohm)) before and after the co-stimulation of the device and followed autoclaving were not significantly different proving that the SMEA device can endure high heat at high pressure during autoclaving and, thus, can be reused. After completing one cycle of the culture and stimulation experiment, we reused the undamaged SMEA devices for one more cycle.

To investigate monitoring of cell viability, the comparison of the ECIS spectra of the electrodes measured at d 1 and d 8 without (**Fig. 2g**, upper panel) and with H9C2 (embryonic rat myoblast) cells (**Fig. 2g**, lower panel) was conducted. The results showed that the device with the cells resulted in increased impedance at a high-frequency range whereas the 17 impedance change of the device with no cells was not significant. In addition, long-term intermittent impedance monitoring of the devices without (upper panel) and with (lower panel) the cells in culture medium was carried out every 2 - 3 h at 10 kHz for 43 h (**Fig. 2h**). In the device with no cells, the impedance was not changed over time. As the cells in the loaded device increasingly adhere to the electrode surface, the impedance value was increased due to the interference of the flow of current by the increased adhesion (stage II, bottom panel of Fig. 2h). Therefore, the impedance value decreased gradually and returned to the state before cell seeding (stage III, bottom panel of Fig. 2h). The factors affecting impedance change include proliferation, cell membranes, adhesion of cells onto electrodes, and gaps between cells^34–36^. The doubling time of the H9C2 cells is 24±5 h^37^, suggesting that there is a step for doubling or movement of the cells at stage III, which weakens the adhesion of the cells on the electrode. This tendency was also shown for other SMEAs (**Supplementary Fig. 4**). Through ECIS measurements, it was possible to confirm not only the presence or absence of the cells but also the degree of cell adhesion to the electrode surface and the degree of bonding between cells. The manufactured device can withstand a long-term cell culture environment and can consistently play multi-functional roles in cell culture, in both monitoring and electro-mechanical co-stimulation.

### Proliferation and differentiation of hiPSCs and co-stimulation of hiPSC-CMs

The overview of the experimental flow for the differentiation of hiPSCs into CMs on the stretchable SMEA device is depicted in **Fig. 3a**. Before the differentiation of hiPSCs, the SMEA device was treated with O_2_ plasma to create a hydrophilic surface which enhances cell adhesion and removes possible organic contaminants^38, 39^. Subsequently, the device surface was treated with recombinant human protein vitronectin (VTN-N), which plays an important role in cell adhesion and migration and eventually promotes cell growth^40^. The cell number density was optimized during the study since confluency plays an essential role in the differentiation of hiPSCs into CMs followed by their maturation^41–43^. On d 0, hiPSCs at a confluency of 4×10^4^ cells/well were seeded onto the SMEA device and allowed to proliferate for 3 days in a large incubator. On d 4, the hiPSCs were treated with CM differentiation factors and allowed to differentiate until d 12. During this period, the hiPSCs were differentiated into CMs and were subsequently subject to co-stimulation. The electro-mechanical co-stimulation was applied for 1 hour every day for 5 days starting from d 13. The viability of the control and stimulated hiPSC-CMs was carried out on d 18 and ECIS from d 0 to d 19.

**Figure 3.**
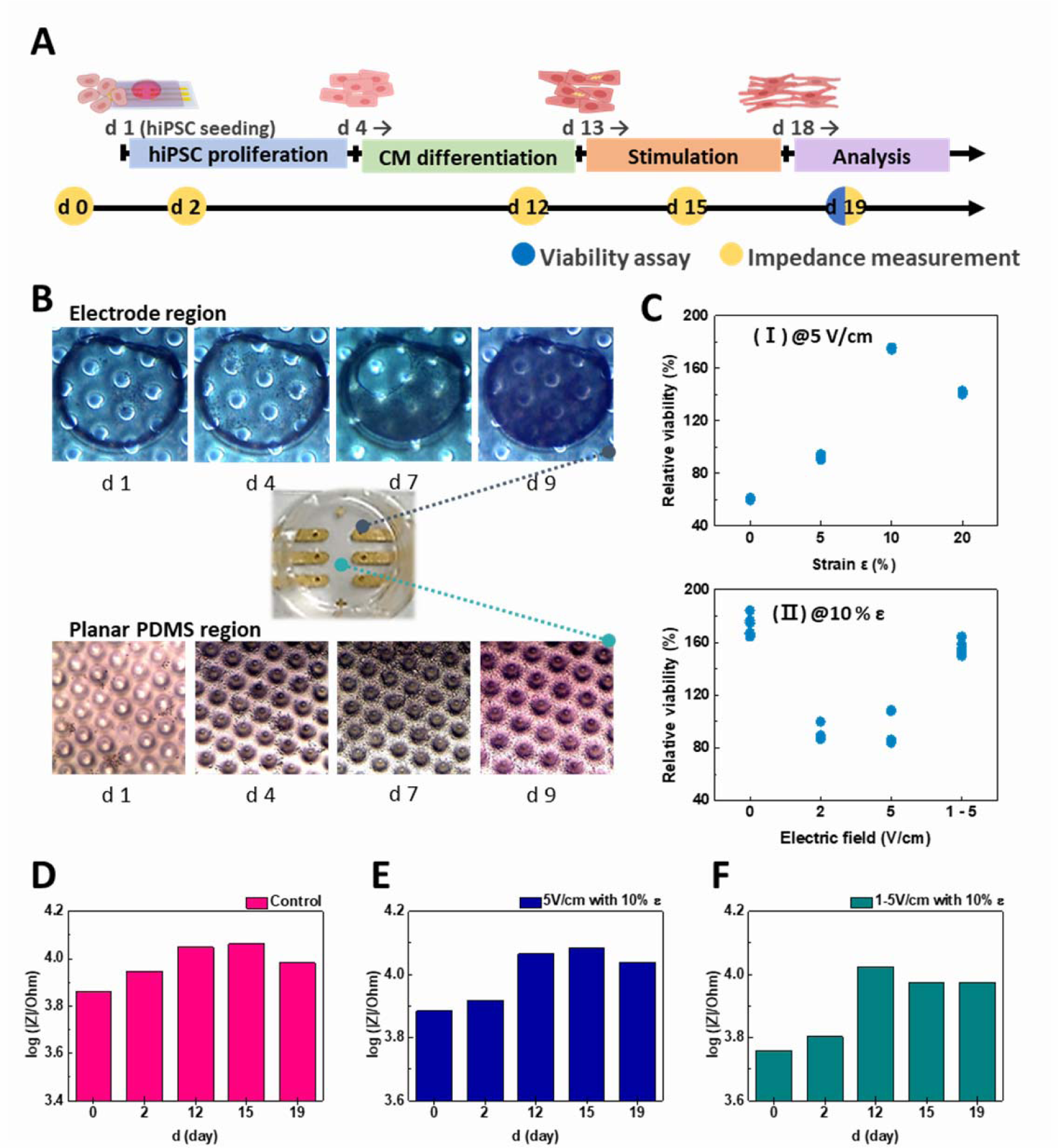
Proliferation and differentiation of hiPSCs and stimulation of hiPSC-CMs. **A**, Experimental flow from hiPSC seeding to analysis. d 1: hiPSC seeding, d 1–d 3: proliferation of hiPSCs, d 4–d 12: differentiation of hiPSCs into CMs, d 13: after differentiation, d 13–d17: co-simulation, d 18: proliferation assay, d 19: 48 h after last simulation. **B**, Optical images of the cells on the electrode area. Seeded hiPSCs are adhered (d 1), proliferated (d 4) and differentiated into CMs (d 7 and d 9). **C**, Relative viability (%) was evaluated by comparing the cell density relative to the control with no stimulations, (I) is the result of various mechanical stimulations with a constant electrical stimulus (E = 5 V/cm) and (II) is the result of various electrical stimulations with a constant mechanical stimulation (ε = 10%). (n=3) **D**, **E**, **F**, Impedance (log(|Z|/Ohm) values of the control, stimulated at 5 V/cm of E and 10% of ε, and at 1-5 V/cm of E and 10% of ε, respectively, for 5 days (d 13–d 18) from the condition before cell seeding to the stage of co-stimulation.

Optical microscopy observations indicated that the hiPSCs seeded onto the electrode areas and planar regions of the encapsulation layer were distributed at d 1 and proliferated and differentiated from d 2 to d 9 (**Fig. 3b**). The optical images obtained on d 4 show that the density of the hiPSCs was increased indicating the active proliferation of the cells on both regions. After differentiation factors were applied, the hiPSCs started to differentiate into CMs (see the optical images obtained on d 7 and d 9). As seen from the optical images, the electrode area and region between the electrodes are highly populated with hiPSCs.

In our initial studies, we aimed to establish a range of optimal electrical and mechanical conditions for co-stimulation, which would keep cell proliferation and viability while minimizing any potential damage to the cells. For co-stimulation of hiPSC-CMs on the SMEA device, uniaxial cyclic stretching and biphasic electrical stimulation with 1 ms of duration time were applied. Both stimulations were applied with a frequency of 1 Hz considering the blood pump mechanism of the heart contraction and relaxation. The normal pulse rate in healthy adults is 60-100 beats per min (1 - 1.6 Hz). By repeating the process of contraction and relaxation, the volume of the heart changes temporarily^44^. For this purpose, the hiPSCs-CMs were divided into two groups. The first group underwent cyclic stretching with different ε values of 0, 5, 10 and 20% under a constant E of 5 V/cm while the other group were stimulated under various E values of 1, 2 and 5 V/cm, with a constant stretching ε of 5%. One group was also used to see the effects of gradually increasing E from 1 to 5 V/cm with an increased step of 1 V/cm with the same ε of 10% during cyclic stretching. After each stimulation experiment, we assessed the viability of the stimulated hiPSC-CMs on d 18 to determine whether electro-mechanical co-stimulation affects apoptosis related to mitochondrial activity in the cells. The effects of the co-stimulation with varying conditions of the ε or E values on the relative viability of hiPSC-CMs were analysed and the data are shown in **Fig. 3c** (I) and (II), respectively. The relative viability (%) is the degree of viability of the stimulated cells in comparison to that of the cells without any stimulation on the SMEA device (100%). The results of varying ε at fixed E of 5 V/cm (Fig. 3c (I)) on the viability of hiPSC-CMs showed lower relative viability than that of the control sample at a low ε range of 0% and 5% but higher relative viability of 175% at an ε of 10%. At cyclic stretching with an ε of 20%, the relative viability was decreased compared to an ε of 10% presumably because of damage to the CMs. The results of the varying electrical stimulations (Fig. 3c (II)) on the relative viability of hiPSC-CMs at a fixed ε of 10% showed that the highest relative viability occurred when electrical stimulation was not applied. As the E increased from 0 to 2 to 5 V/cm, we observed a decrease in the relative viability. However, when we increased the E gradually from 1 to 5 V/cm at a fixed ε of 10%, the relative viability increased to over 100%, indicating a level of high viability closer to that of the sample with only mechanical stimulation. This may be attributed to the remodelling of CMs, which allows them to adapt to the electrically stimulated environment. The results show the beneficial effects of co-stimulation, for example, with an ε of 10% and gradually increasing E from 0 to 5 V/cm, on the viability of hiPSC-CMs.

To confirm the capability of the developed SMEA device to monitor the status of the cells continuously during the proliferation of hiPSCs and co-stimulation of hiPSC-CMs. The ECIS was monitored by using the SMEA array as the working electrode, Ag/AgCl electrode as a reference electrode and Pt as a counter electrode. The impedance spectra of the control sample (hiPSCs-CMs without stimulation), the co-stimulated sample at the E of 5 V/cm with ε of 10%, and the co-stimulated sample at a gradual E of 1-5 V/cm with ε of 10% are shown in **Fig. 3d**, **3e** and **3f**, respectively. The impedance values (log(|Z|/Ohm) were measured for hiPSCs at d 0 and d 2, the differentiated hiPSC-CMs before stimulation at d 12, and the stimulated hiPSC-CMs at d 15 and d 19. The impedance value was initially measured for the bare electrode as a baseline with no cell seeding. As shown in **Fig. 3d**, the impedance value of the control sample without any stimulation increased gradually as the hiPSCs proliferate and differentiate (d 2 and d 12), reached a maximum value of 4.06 on d 15 and then decreased slightly on d 19. A similar trend was observed when the hiPSC-CMs were co-stimulated with an E of 5 V/cm and ε of 10%, the impedance value reached a maximum value of 4.08 on d 15 and then decreased slightly after co-stimulation (**Fig. 3e**). For the sample with a gradual increment in the E under a constant ε of 10%, the impedance value reaches its maximum at the end of differentiation on d 12 and decreased slightly during and after co-stimulation (**Fig. 3f**). The observed trends on the ECIS measurements from d 0 to d 19 are attributed to the changes in the number, movement and adhesion of the cells on the electrode during proliferation, differentiation and co-stimulation. The hiPSCs initially proliferate to reach a certain confluence, and then differentiation begins by terminating further proliferation of hiPSCs resulting in the increase of the impedance values from d 2 and d 12. From d 13 to d 19, the proliferation of hiPSC-CMs almost does not occur but the cell-to-cell contraction, fusion, migration and maturation of hiPSC-CMs would have primarily occurred resulting in the change of impedance values. When the hiPSC-CMs were investigated after co-stimulation, the building of cell bundles resulted in the formation of a sheet of hiPSC-CMs due to the close intercellular communication among the differentiated CMs (**Supplementary Fig. 5**). The formation of cell sheet tissue on the electrodes resulted in an increase in the open areas on the electrodes as a result of cell-to-cell binding, which may contribute to a decrease in the impedance values. Since the ECIS is a label-free non-invasive assay to measure intercellular junction and cell coverage of working electrodes, the ECIS of the cells on the SMEA in the multi-functional tissue engineering system can provide indirect information on the viability of cells that are present on the electrodes. In addition, the capability of ECIS on the SMEA device provides a useful tool for monitoring the different stages of proliferation, differentiation and stimulation in line^34–36^.

### Effects of co-stimulation on the maturation of hiPSC-CMs

We further analysed the effects of various combinations of co-stimulation conditions on the maturation degree of hiPSCs-CMs. For this purpose, the hiPSC-CMS were labelled with fluorescently tagged antibodies for ICC analysis. Two key markers, NKX2-5 which signifies the early cardiac mesoderm and TNNT2 which is an important marker of myocardial cells, were used to study the effect of co-stimulations on the maturation of hiPSC-CMs. **Fig. 4a** shows the confocal microscopy images of the co-stimulated hiPSCs-CMs when the E value was kept at a constant 5 V/cm and the ε was varied. **Fig. 4b** represents the immunostaining results of hiPSCs-CMs stimulated at a constant ε of 10% while the E value was varied. With or without applied electrical stimulation, when an ε of 10 or 20% was applied, the stimulated hiPSC-CMs are aligned with increased directionality (white arrows in Fig. 4). The alignment characteristics of the hiPSC-CMs were analysed statistically using the images of TNNT2 expression in Fig. 4. It is evident from the comparison between the control and stimulated samples that there are clear differences in the directionality of the stimulated samples (**Supplementary Fig. 6**). From the expression of TNNT2 in Fig. 4a, the fibre bundles of CMs in the samples with the ε of 5% and 10% were more prominent compared to the control sample with no stimulation, and the fibre bundles of CMs in the sample with an ε of 20% were less significant compared to the sample stimulated with an ε of 5 % and 10 %. The stronger the mechanical stimulation at a constant E of 5 V/cm, the more directional the fibre bundles. However, the maturity observed from the fibrous structure of TNNT2 expression suggests that proper mechanical stimulation is necessary, given the degree of subdivision in the fibrous structure. When CMs are matured, the expression of TNNT2 increases and the proteins become more organized into a fibrous structure^13, 45^. When the results of the relative viability (Fig. 3c (I)) and ICC (Fig. 4a) are considered, it can be assumed that the ε of 10% at the E of fixed 5 V/cm is a balanced condition for both cell viability and maturation. When the E was varied at the fixed ε, it was difficult to confirm the significant difference in the TNNT2 fluorescence expression intensity between the control sample and the samples with the E of 0 and 2 V/cm (Fig. 4b). However, the ICC images of hiPSC-CMs stimulated with the E of 5 V/cm and a gradual E of 1-5 V/cm at a fixed ε of 10% indicate that the expression intensity of TNNT2 is stronger and clearer at higher E and thin and dense fibres are well distributed compared to the control sample (Fig. 4b). The higher relative viability of the samples co-stimulated with an E of 5 V/cm and 1-5 V/cm at a fixed ε of 10% than the samples co-stimulated at lower E (Fig. 3c (II)) were qualitatively consistent with the ICC data in Fig. 4b.

**Figure 4.**
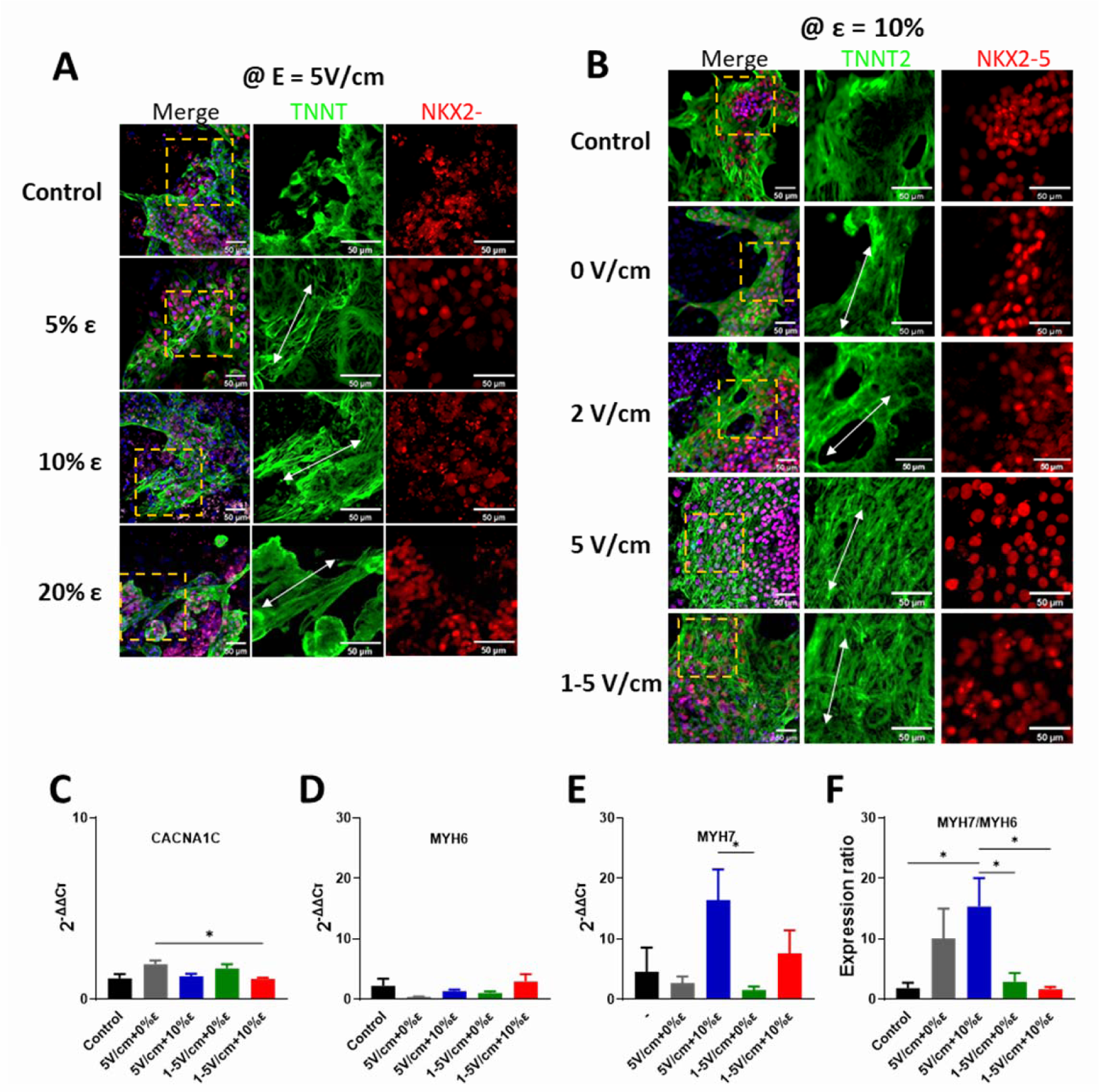
Analysis of the effects of stimulated mechanical and electrical co-stimulation on cell differentiation and maturation by ICC and qRT-PCR. **A**, ICC images of the samples with various ε at a constant E of 5 V/cm, **B**, ICC images of the samples with various E value at a constant ε of 10%. Scale bar is 50 μm. **C-F**, Gene expression analysis data of non-stimulated control hiPSC-CM sample, stimulated hiPSC-CMs with an E of 5 V/cm and ε of 0% and 10%, and gradual E of 1-5 V/cm at the ε of 0 and 10%. CACNA1C: alpha-1 subunit of a voltage-dependent calcium channel, MYH6: myosin heavy chain alpha isoform, MYH7: myosin heavy chain beta isoform, *Statistically significant differences between control, 5 V/cm with 0% ε, 5 V/cm with 10% ε, 1-5 V/cm with 0% ε and 1-5 V/cm with 10% ε. Error bars represent the mean value ± SEM, the sample number of control and 5 V/cm with 0% ε for MYH7 = 2, the sample number of stimulated samples except the one with 5 V/cm at 0% ε for MYH7 = 3. Every sample was analyzed three times repetitively. p < 0.05, one-way ANOVA with post hoc Tukey test.

The fluorescence expression of NKX2-5, which represents early cardiac mesoderm, did not show a clear tendency while intensity was maintained throughout the co-stimulations (Fig. 4a and 4b). It has been reported that the expression of NKX2-5 is decreased in long-term cultures^46^. The observed expression intensity was kept presumably because the period of the culture was not very long. Since the ICC data are qualitative, the expression data of NKX2-5 and TNNT2 were not enough to reveal all their features at every region of the samples and so it was difficult to fully judge the effects of co-simulation conditions on the maturation of hiPSC-CMs with ICC alone.

To complement the limits of ICC analysis, for example, higher viability of the sample stimulated at the E of 0 V/cm with ε of 10% but lower maturation than other co-stimulated samples, quantitative reverse transcription polymerase chain reaction (qRT-PCR) analyses were performed. The qRT-PCR was used to investigate the effects of co-stimulation on the expression of various genes for the samples with salient features in the viability assays (Fig. 3c) and ICC analysis (Fig. 4). A compilation of the gene expression in various co-stimulated hiPSC-CMs is represented in **Fig. 4c-4f**. The expression of the *CACNA1C* gene is related to the electrophysiological characteristics of action potential duration and calcium handling of storage, cycling and dynamics. As shown in **Fig. 4c**, the difference in the gene expression of *CACNA1C* was not significant for the co-stimulated samples compared to the control samples, indicating that maturity in their electrical properties was not clearly observed. A comparison of *MYH6* (encodes myosin heavy chain alpha (MHC-α) isoform) and *MYH7* (encodes myosin heavy chain beta (MHC-β) isoform) gene expression in samples under different stimulation conditions is shown in **Fig. 4d** and **4e**, respectively. In Fig. 4d, samples stimulated with a gradual E of 1-5 V/cm at a fixed ε of 10% expressed the highest level of the *MYH6* gene (2.98±1.12), followed by the control and the sample co-stimulated with an E of 5 V/cm at an ε of 10% (2.27±1.06 and 1.36±0.25, respectively). The data in **Fig. 4e** shows that a much higher gene expression level was observed in *MYH7* (16.44±4.76 and 7.65±3.55, respectively) especially when an ε of 10% was applied to the hiPSC-CMs on the SMEA devices with an E of 5 V/cm and gradual E of 1-5 V/cm, respectively. As hiPSC-CMs mature, the MHC-α isoform shifts to the MHC-β isoform^2, 13, 24, 47, 48^ and, in turn, the expression ratio of *MYH7*/*MYH6* is increased with an increase in the maturation of hiPSC-CMs^2, 49^. In the expression ratio of *MYH7*/MYH6 plotted using the data in Fig. 4d and 4e, the *MYH7*/*MYH6* expression ratios of the stimulated hiPSC-CMs were higher than that of the control sample (**Fig. 4f**). The sample co-stimulated with an E of 5 V/cm at an ε of 10% expressed the highest *MYH7/MYH6* gene expression ratio (15.33±4.46), which indicates the highest degree of maturation. The results confirm that the co-stimulation enhanced the degree of maturation in hiPSC-CMs.

### Role of co-stimulation on calcium transient and membrane potential

Fluorescent signals of the calcium transients from the hiPSC-CM samples of the control without any stimulation and co-stimulated with an E of 5 V/cm and a gradual E of 1-5 V/cm at an ε of 10% were imaged using fluorescent confocal imaging to investigate whether co-stimulation affects the characteristics of calcium transients at d 19. For hiPSC-CMs, a time-dependent transitory increase in intracellular calcium ions (Ca^2+^) plays a significant role in the contraction. This change in the Ca^2+^ concentration is prompted by the membrane potential that spreads throughout the heart. Hence, detecting the spatiotemporal dynamics of Ca^2+^ is vital to confirming the maturation of a cardiac tissue construct. Hence, we used a Ca^2+^-sensitive dye as an indicator to confirm intracellular calcium signalling. The results of gene analysis with CACNA1C, which indicates electrophysiological characteristics, were difficult to compare in different stimulation conditions. The calcium transient analysis can complement the ICC analysis. Previous studies have suggested that an increment in the amplitude of the Ca^2+^ influx and a decrease in the peak width denote the differentiation of hiPSCs to CMs and the maturation of hiPSC-CMs^13, 15, 50^. The fluorescent confocal images of the calcium transients indicate that all of the hiPSC-CMs have excellent intracellular calcium signalling functions (**Fig. 5a**). However, it is also evident from the images that hiPSC-CMs were better aligned with simultaneous electro-mechanical co-stimulation (Fig. 5a(I) and (II)) compared to the control sample (Fig. 5a(I)). For quantitative analysis, the amplitude, SlopeMax2Pk value in the y-axis, which is the absolute maximum slope from peak start to peak maximum, and full width at half-maximum (FWHM) values were analysed using the recorded calcium transients for the control and stimulated samples (**Supplementary Videos 1-6) and** are shown in **Fig. 5b, 5c** and **5d**, respectively. The amplitude values for the control and co-stimulated hiPSC-CMs clearly indicate that the simultaneous electro-mechanical co-stimulation modulated the amplitude of the signals more significantly compared to that of the control sample (Fig. 5b). The SlopeMax2Pk data obtained indicated significantly higher SlopeMax2Pk value for the co-stimulated samples when the E value was gradually increased from 1 to 5 V/cm at the fixed ε of 10%, compared to the control sample (Fig. 5c). It is evident from the analysed results, as shown in Fig. 5d, that the hiPSC-CMs co-stimulated with a gradual E of 1-5 V/cm at a constant 10% of ε showed the narrowest FWHM value indicating enhanced maturation characteristics of hiPSC-CMs compared to the control sample or the sample co-stimulated with an E of 5 V/cm at an ε of 10%. The obtained results are consistent with results reported previously^13, 15, 50^, the matured hiPSC-CMs are characterized by increased values of the signal amplitude and SlopeMax2Pk and reduced FWHM values compared to unmatured hiPSC-CMs. Overall, our analysed results of the gene expression and calcium transients indicate that the maturation of hiPSC-CMs was more prominent when an electrical stimulation with an E of 5 and 1-5 V/cm at an ε of 10% was applied.

**Figure 5.**
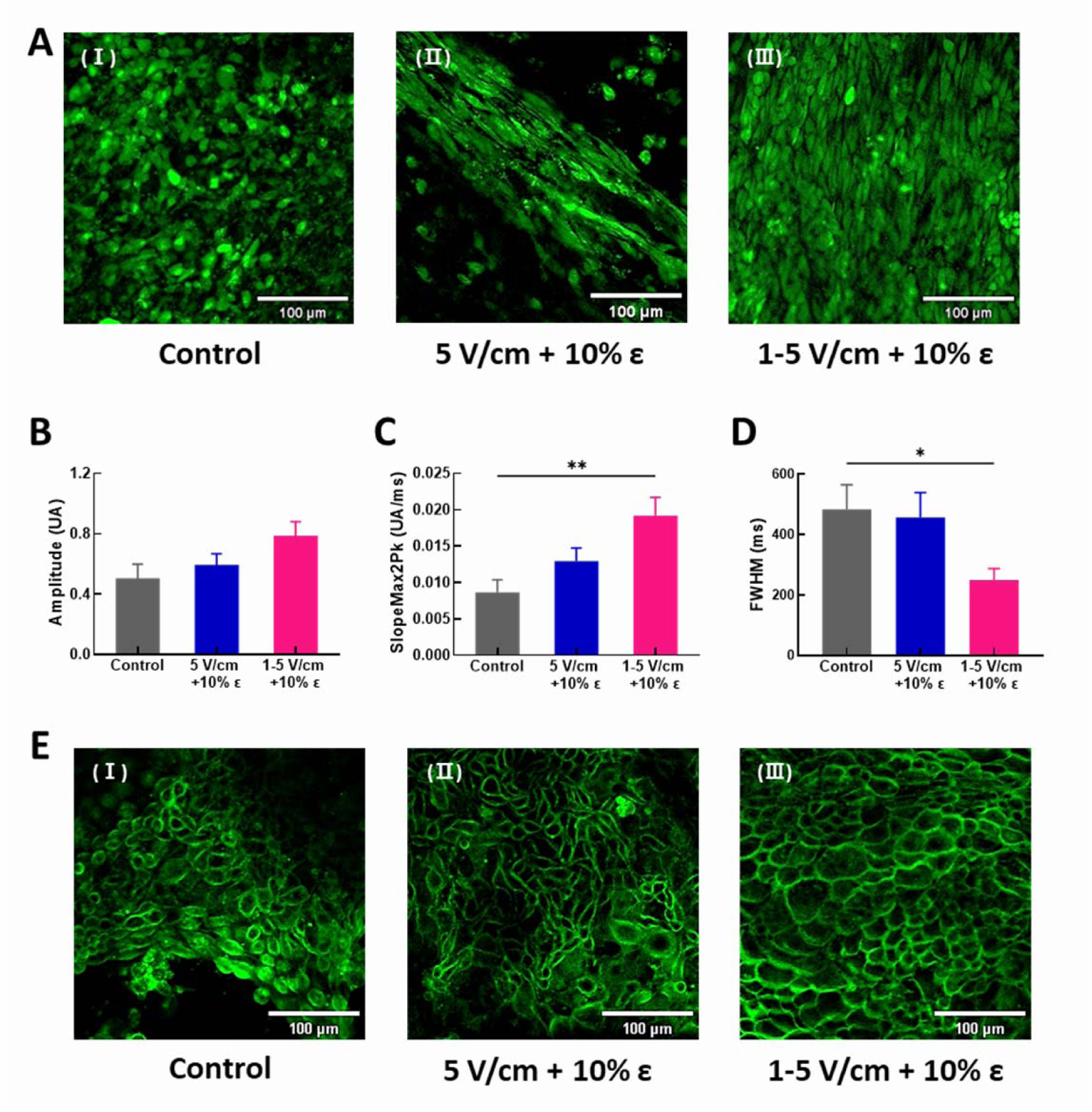
Analysis of calcium transient and membrane potential. **A**, Fluorescent confocal images of calcium transients of the control hiPSC-CMs (I) and of the hiPSC-CMs co-stimulated with the E of 5 V/cm (II) and the gradual E of 1-5 V/cm (III) at an ε of 10%. The statistical data of **B**) amplitude, **C**) absolute maximum slope from peak start to peak maximum (slopemax2Pk, and **D**) full width at half maximum (FWHM) obtained from the calcium transients of hiPSC-CMs (the number of samples, n = 1 per group, the number of measuring points = 9-10 points, exposure time = 50 ms, p× < 0.05, ** < 0.005 one-way ANOVA with post hoc Tukey test) **E**, Fluorescent confocal images of membrane potential of the control hiPSC-CMs (I) and of the hiPSC-CMs co-stimulated with an E of 5 V/cm (II) and a gradual E of 1-5 V/cm (III) at the ε of 10%.

Alterations in the electrical potential across the membrane of hiPSC-CMs also have significant implications in various physiological parameters such as cell signalling and the contraction of cardiac muscle fibres. The analysis of electrical potential and conduction in hiPSC-CMs is important since signal propagation symbolizes a healthy cardiac construct^15, 51^. Therefore, the membrane potential was measured on d 19 to determine how co-stimulation can affect the maturation of the differentiated hiPSC-CMs. We conducted voltage imaging of hiPSC-CMs using a membrane-sensitive fluorescent dye and confocal microscopy. Confocal fluorescent images were obtained from the control without any stimulation and hiPSC-CMs co-stimulated with an E of 5 V/cm and 1-5 V/cm at a fixed ε of 10%. The fluorescent images revealed excellent electrical membrane functions from all the hiPSC-CMs (**Fig. 5e**). The results demonstrate that hiPSCs were effectively differentiated into CMs, exhibiting a consistent electrical function related to membrane potential, regardless of the presence or absence of stimulation.

## 4. Discussion

The adult human heart has an extremely low regeneration rate, less than 1% per year, which makes self-recovery difficult when heart cells are damaged by conditions such as myocardial infarction^52^. Moreover, using cardiac muscle cell-based treatments has serious limitations, for instance, when adult human heart cells are extracted they lose the characteristics of intact adult heart tissues, limiting their use for treating various cardiac diseases^53, 54^. To overcome these limitations, the differentiation of SCs into CMs, including embryonic SCs^25, 55–57^ and hiPSCs^24–26, 46, 58^, has been investigated extensively over the last decade. The use of hiPSCs for cardiac tissue engineering is particularly promising because there are no ethical issues. However, hiPSC-CMs have maturation issues^11–13, 15, 16^ because they resemble foetal-state CMs and have not gone through the process of maturation with years of mechanical and electrical stimulations like the human heart after birth^5^. Therefore, for applications involving hiPSC-CMs, the development of approaches to induce their maturation is essential. Previous studies on the simultaneous application of multiple stimuli to hiPSC-CMs have reported synergistic effects and a higher level of maturation^59^. However, the detailed maturation mechanism with multiple external stimuli has not been identified, and conclusive outcomes are still lacking under different stimulation conditions. Furthermore, previous research on co-stimulation systems has limits in the hardware for long-term culture with stimulation and constraints. For example, they provided cyclic mechanical stretching stimulation, but a completely integrated electrical and mechanical co-stimulation system could not be established because structurally stable and durable electrodes are difficult to be formed monolithically on stretchable substrate^28, 60, 61^. Alternatively, mechanically static stimulation rather than dynamic stimulation is often used even though electrical stimulation is sufficiently applied^20, 23, 24, 29^. There is a dearth in the variety of co-stimulation approaches as a result of the limitations of many technologies.

Our approach to obtain matured hiPSC-CMs by applying various electro-mechanical co-stimulation conditions that simulate the growth and maturation of the CMs is promising. In developing an *in vitro* multi-functional tissue engineering system, we prioritized the fabrication of highly durable electrodes for long-term monitoring and co-stimulation in culture media. The approach could be realized by creating a highly durable SMEA array device with a culture well as a key component. This device enabled the proliferation, differentiation, and electro-mechanical co-stimulation of hiPSCs and hiPSC-CMs by forming the electrodes on a stress-absorbing 3D-patterned stretchable substrate and encapsulation. The SMEA device was also transparent, allowing the optical microscopy of hiPSCs and hiPSC-CMs without damaging the devices or cells. The monolithic integration of multiple components and dual function of the multi-electrodes on the same substrate allowed a simple and compact structure of the SMEA device and a small footprint so that it can be integrated into a mini-incubator. This advantage allowed the cells to be kept outside of a large incubator during co-stimulation and monitoring. Additionally, the excellent stability of the device and durability against electrical and mechanical stimulations in culture media allow long-term culture and autoclaving for reuse. The ECIS of cells on the SMEA array electrodes could provide useful information on the cell status during stages of proliferation and differentiation of hiPSCs and co-stimulation of differentiated CMs.

Using the multi-functional tissue engineering system, various electro-mechanical co-stimulations were applied to hiPSC-CMs in search of optimal conditions for their maturation. By controlling co-stimulation parameters, the shape and arrangement of CM fibres and their maturity level could be manipulated. Maturity was confirmed through ICC and gene expression analyses. Additionally, analysis of calcium transients and membrane potential measurements showed that the electrical properties of co-stimulated hiPSC-CMs were improved. The simultaneous application of cyclic mechanical deformations of 10% and gradually increasing electrical field resulted in hiPSC-CMs with highly desirable electrical properties.

In summary, our multi-functional engineering system demonstrated the ability to perform electro-mechanical co-stimulation with an independent control of electrical and mechanical stimuli, as well as continuous monitoring of cells using optical and electrochemical methods, without significant damage to the cells or devices. This system has the potential to be a valuable tool in the field of regenerative medicine for electrogenic cells including CMs, skeletal muscle and neuronal cells, specifically in the use of hiPSCs. By integrating the SMEA device with stimulation control systems, a wider range of experimental investigations can be conducted beyond the scope of this study. In addition, mechanical cues associated with scaffolds and chemical cues can be added without difficulty. This approach has promising potential for tissue engineering of electrogenic cells, drug development, drug toxicity evaluation and patient-specific drug and disease research using hiPSC-CMs.

## Acknowledgments

This work was supported by the Nanomaterial Technology Development Program (No. 2021M3H4A4079519) and the Basic Science Research Program (No. 2019R1A6A1A03033215) through the National Research Foundation of Korea (NRF) funded by the Ministry of Science, ICT, & Future Planning and the Ministry of Education. We thank Dr Le Thai Duy for helping to draw the schematics of the device and system.

## Author Contributions

A.-R.K. carried out the experiments, analysed the data and wrote the manuscript. S.S. discussed the ideas and reviewed and edited the manuscript. H.B.L. designed and provided the 3D micro-patterned PDMS substrate master mould. N.-E.L. planned and supervised the work and reviewed and edited the manuscript.

## Novelty and Significance

What is it known?

- The utilization of cardiomyocytes derived from human induced pluripotent stem cell (hiPSC-CMs) presents an exceptional chance to model cardiac diseases specific to patients, explore drug discovery, conduct screening for cardiotoxicity, and develop therapy using stem cells.
- The existing methods for differentiating hiPSCs into cardiomyocytes have not been able to fully replicate the electro-mechanical function of the myocardium, which restricts their potential applications.
- Encouraging advancements have been made in enhancing the maturation of hiPSC-CMs through the use of mechanical and electrical stimulations. However, optimizing multiple factors is essential to achieve the most effective outcomes for consistent long-term culture and stimulation.

What new information does this article contribute?

- Our approach of utilizing diverse electro-mechanical co-stimulation conditions that mimic the development and maturation of cardiac constructs shows great potential in achieving matured hiPSC-CMs.
- Our developed SMEA device has a simple and compact structure with a small footprint, thanks to the integration of multiple components and dual functions. This enables its integration into a mini-incubator, facilitating co-stimulation and monitoring of cells outside of a large incubator.
- The device’s outstanding stability and durability to both electrical and mechanical stimulation in culture media permit long-term culture and autoclaving for reuse.

The utilization of hiPSC-CMs has provided a unique avenue for modeling patient-specific cardiac diseases, developing stem cell-derived cardiac tissue therapies, exploring drug discovery, and conducting cardiotoxicity screening. However, current differentiation procedures have yet to fully replicate the electro-mechanical function of the myocardium, which limits their potential applications. Exciting advancements have been made in enhancing the maturation of hiPSC-CMs by employing diverse electro-mechanical co-stimulation conditions that mimic natural development, showing great promise in achieving matured cardiomyocytes (CMs). A notable solution is the SMEA device, which boasts a simple and compact structure that can be seamlessly integrated into a mini-incubator. This integration enables co-stimulation and monitoring of cells outside of a large incubator, overcoming previous constraints. Furthermore, the SMEA device exhibits exceptional stability and durability when subjected to electrical and mechanical stimulation in culture media, enabling long-term culture and autoclaving for reuse. Consequently, the multi-functional tissue engineering platform based on the SMEA device offers an excellent option for experiments requiring prolonged culture, monitoring, differentiation, and maturation.

